# A Self-Supervised Machine Learning Approach for Objective Live Cell Segmentation and Analysis

**DOI:** 10.1101/2021.01.07.425773

**Authors:** Michael C. Robitaille, Jeff M. Byers, Joseph A. Christodoulides, Marc P. Raphael

## Abstract

Machine learning algorithms hold the promise of greatly improving live cell image analysis by way of (1) analyzing far more imagery than can be achieved by more traditional manual approaches and (2) by eliminating the subjective nature of researchers and diagnosticians selecting the cells or cell features to be included in the analyzed data set. Currently, however, even the most sophisticated model based or machine learning algorithms require user supervision, meaning the subjectivity problem is not removed but rather incorporated into the algorithm’s initial training steps and then repeatedly applied to the imagery. To address this roadblock, we have developed a self-supervised machine learning algorithm that recursively trains itself directly from the live cell imagery data, thus providing objective segmentation and quantification. The approach incorporates an optical flow algorithm component to self-label cell and background pixels for training, followed by the extraction of additional feature vectors for the automated generation of a cell/background classification model. Because it is self-trained, the software has no user-adjustable parameters and does not require curated training imagery. The algorithm was applied to automatically segment cells from their background for a variety of cell types and five commonly used imaging modalities - fluorescence, phase contrast, differential interference contrast (DIC), transmitted light and interference reflection microscopy (IRM). The approach is broadly applicable in that it enables completely automated cell segmentation for long-term live cell phenotyping applications, regardless of the input imagery’s optical modality, magnification or cell type.

## Introduction

Live cell phenotyping is an information rich experimental approach, capable of providing mechanistic insights into cell biology^1,2^, guiding drug development^3^ and elucidating disease pathologies^4,5^. The wealth of information available from live cell microscopy results from the fact that there are numerous optical modalities that can be integrated within a given experiment – from fluorescence imaging which provides spatio-temporal information on specific signaling pathways and organelles to label-free techniques such as phase contrast and differential interference contrast (DIC) which enable the visualization of whole cellular morphologies and dynamics. Each of these modalities provides its own outcome measures which can be viewed as static snapshots or dynamic variations within the four-dimensional space of x, y, z and time^6^.

However, compared to genotyping - its synergistic partner technique - live cell phenotyping remains a far more subjective science. The generation of genomic sequencing data and its analysis can now be achieved autonomously by employing a combination of robotics and microfluidics for sample preparation and machine learning algorithms for data collection and interpretation. In contrast, the extraction of quantitative information from live cell imagery by manual means is still commonplace in live cell microscopy, a fact which speaks to the human visual system’s adeptness at detecting small changes and low contrast features with high fidelity. But with automated live cell microscopes now able to collect high resolution imagery for days on end, the resulting data files can quickly grow to tens of gigabytes, leaving the analyst with an overwhelming amount of imagery to work through.^7^ Furthermore, if the analyst is not blinded to the experimental design, unconscious bias can creep into the data extraction process.

Enter computational algorithms capable of extracting the relevant outcome variables from the imagery in an automated fashion.^8-10^ Broadly speaking, the algorithms are often classified as model based approaches (e.g. Cell Profiler)^11^, and machine learning algorithms, (e.g. U-Net, ilastik)^12-14^. Neither approach is completely autonomous when it comes to cell segmentation: model-based approaches require the manual tuning of multiple parameters, while machine learning requires the user provide curated data from which the algorithm is trained. Once tuned or trained, the software is able to process far more imagery than could be achieved manually - but there is still a human-in-the-loop. It is just that the manual contribution has been moved to the front end for training purposes and is then continuously reapplied by the algorithm. Algorithms that are tuned or trained at the onset can problematically miss relevant features as the cellular phenotypes or background characteristics evolve, inadvertently skewing the analysis. For instance, variations in label intensity (e.g. photobleaching, quenching) or new morphological features that were not present during the initial training (e.g. differentiation, mitosis, blebbing) can go undetected if not retrained with a freshly curated data set or parameters that capture the offending features. In the same way, temporal variations in the background illumination intensity or homogeneity can also result in improper cell segmentation.^7^ Especially concerning is that the user-supervised training process is inherently subjective in nature and can cause unconscious biases to be effectively baked in to the extracted data by the training process. To optimize objectivity and efficiency, an essential goal is to develop software that can accept imagery from any optical modality, labeled or unlabeled, and extract the cellular features of interest *without input from the user*.

As participants in a synthetic biology real-time reproducibility project administered by U.S. Defense Advanced Research Projects Agency (DARPA), referred to as Independent Verification & Validation (IV&V), we have recently experienced all of these algorithmic limitations and how they can result in large amounts of data either being incorrectly segmented, subjectively segmented, or left unanalyzed due to time constraints.^15^ The program involves a wide range of cell types (amoeboid to eukaryotic) from multiple cell biology laboratories; multiple imaging modalities – both fluorescent and label free; and objective magnifications ranging from 10X to 100X. The cumbersome process of retraining supervised machine learning software to match this variety of conditions proved impractical and a human-in-the loop training step was deemed too subjective. The challenge then was to develop a completely automated segmentation algorithm for live cell microscopy applications. In particular, the image analysis software should be ‘self-supervised’, meaning it trains itself to classify cells versus background and then regularly updates this training so that it can adapt to evolving intensities and morphologies. The software was required to segment a variety of cell types from live cell imagery given the most common imaging modalities as inputs - phase contrast, transmitted light, DIC, fluorescence and interference reflection microscopy (IRM) – and to do so without user-adjustable parameters or user-selected training imagery. It was additionally required that the generated models adapt to changing cell phenotypes and lighting conditions for long-term imaging applications (hours to days).

## Methods

To replace more manual model based and machine learning training approaches for segmenting cells with an automated, self-supervised algorithm, we took advantage of the one phenotypic feature which is present in live cell microscopy no matter what the modality: *motion*. From the nanoscale diffusion of proteins and vesicles to the migration of cells that are tens of microns in length, the ever present dynamics captured by live cell microscopy make it ideal for applying optical flow (OF) algorithms designed to identify not just spatial intensity features in a given frame but also the variation or ‘flow’ of those features from frame to frame. The central assumption in optical flow algorithms is that the overall image intensity will remain constant if the time difference between frames is reasonably small.^16^ This leads to the following time-derivative constraint equation:

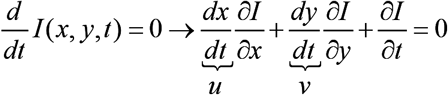

where *I*(*x, y, t*) is the in-plane image intensity at time *t, u* and *v* being the optical flow in the *x* and *y* directions, respectively. The methods used to solve this constraint equation are matched with the imaging goal, such as reducing jitter in imagery taken from helicopters, aligning medical imagery or, in the case of this study, cell motion segmentation. In testing a range of optical flow algorithms for cell segmentation, we found the Farnebäck method to be the most robust due to its sensitivity to object deformation – a natural fit for cells which are morphologically variable.^17,18^

OF assumptions may or may not be met for fluorescence time-lapse imagery applications in which extended time intervals are sometimes employed to avoid phototoxicity or photobleaching.^6,19^ For this reason, it was important that our technique be co-validated with label free techniques such as transmitted light and phase contrast which are minimally invasive. Overlays of less frequently accumulated fluorescence imagery with cells segmented using a label-free imaging channel is then straightforward. Furthermore, there has been an increased appreciation for the morphological information label-free approaches can provide as a result of algorithmic-based phenotyping.^20-22^

Our approach to self-supervised learning and automated model generation begins with utilizing the Farnebäck OF method as a means of classification bootstrapping (Fig 1). Typical segmentation strategies involve utilizing static information in a single image at time frame (*t*), which can have difficulty distinguishing ‘cell’ from ‘background’ pixels in a generalizable manner (Fig 1a). In contrast, our approach begins with an OF calculation based on images from consecutive time frames (*t*-1, *t*). This enables us to leverage the ubiquitous nature of intracellular motion and build a dynamics-based feature vector: pixels with the highest flow are automatically labeled as ‘cell’ pixels, those with the lowest flow are automatically labeled as ‘background’ pixels, and those that do not fit either category remain unlabeled (Fig 1b,c). We note that this automatic self-labeling is broadly applicable in that it is not dependent on principles of any specific optical modality, cell type, or phenotype.

**Fig 1.**
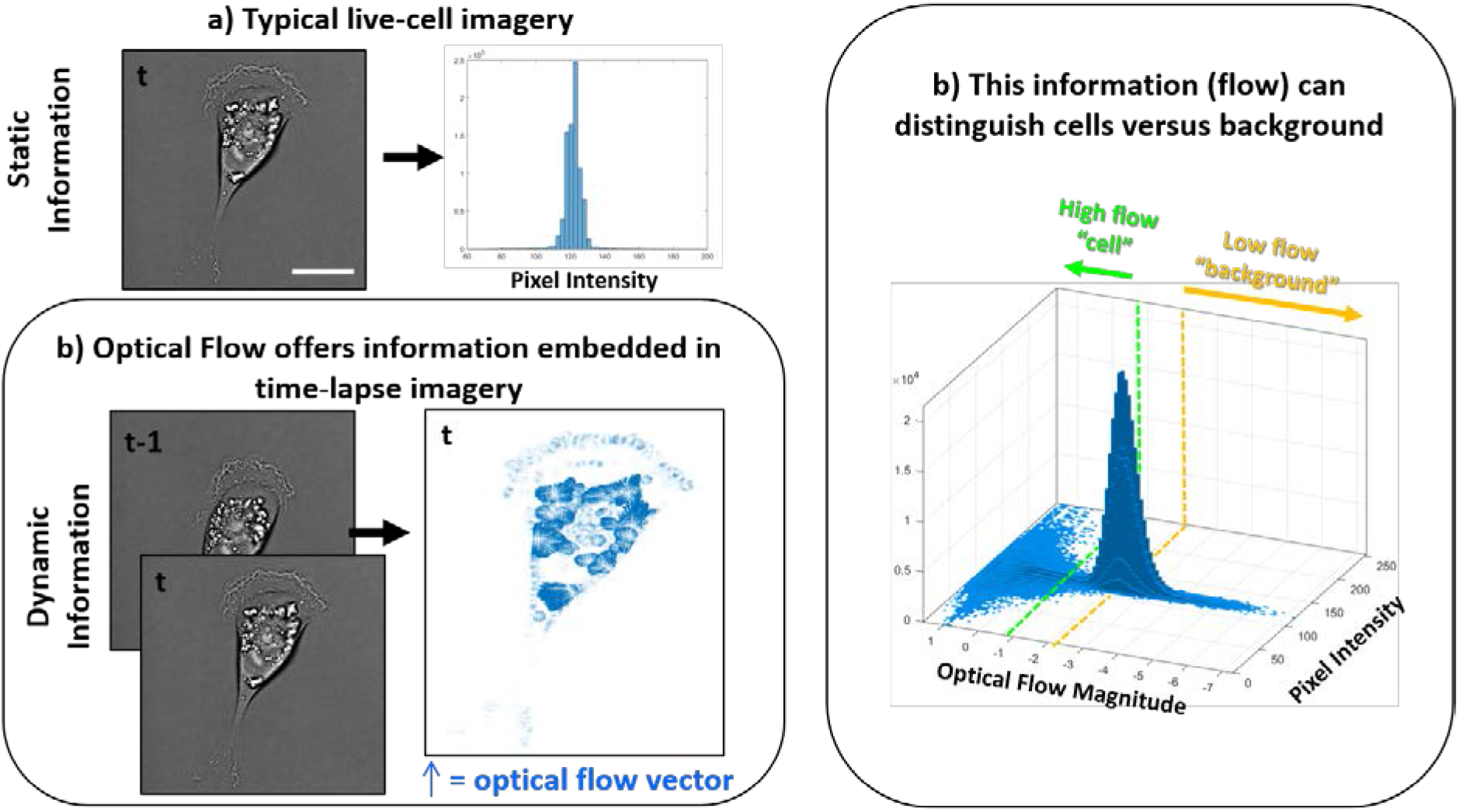
Overview of the optical flow self-labeling strategy. **(a)** The vast majority of cell segmentation techniques utilize single image frames and the static information contained within as means to distinguish ‘cell’ from ‘background’, oftentimes represented in a histogram. The self-supervised algorithm utilizes optical flow as a means to self-label pixels in an automated fashion. **(b)** Due to the prevalence of intracellular dynamics in time-lapse live cell imagery, optical flow can be calculated for each pair of consecutive images (*t* − 1, *t*). The optical flow can then be represented as vectors associated with each pixel (right). **(c)** The magnitude of the optical flow then offers a means to distinguish cells from their background, as shown in the bivariate histogram which co-plots the pixel intensity of a single image at *t* to the optical flow vector magnitudes calculated between consecutive images (*t* − 1, *t*). Pixels with the highest flow can be automatically labeled ‘cell’ (left of the green dashed line) and those with the lowest flow can be labeled ‘background’ (right of the yellow dashed line). Pixels that do not meet either criteria remain unlabeled, while the self-labeled pixels are used to create a training data set for classification. Time increment: 600 sec, scale bar = 20 µm.

The OF-based self-labeling approach outputs a set of ‘cell’ and ‘background’ labeled pixels which are then used to generate additional entropy and gradient feature vectors at each time point. These static feature vectors are used to train and generate a classifier model which, in the final step, is applied to all pixels in the image for cell segmentation. The algorithm is written in stand-alone MATLAB script and utilizes functions from the Image Processing, Statistics and Machine Learning, and Computer Vision Toolboxes.

The self-supervised training approach is illustrated in Fig 2 using time lapse DIC imagery of multiple (top) and a single highlighted (bottom) MDA-MB-231 cell. From the raw imagery (Fig 2a,b), many portions of individual cells appear to blend in with the background. However, when the OF self-labeling strategy is applied, the algorithm automatically identifies pixels with high flow magnitude, highlighted as green pixels (Fig 2c,d), which are selected as having the highest probability of correctly being labeled ‘cell’. To automatically label the background, the algorithm over segments, that is, a liberal (low) OF threshold is employed which captures motion from not only the cell but also from nearby background pixels as well. The algorithm sets these pixel values to zero and labels the pixels in which no significant motion was detected as ‘background’ (Fig 2c,d yellow pixels). Once labeled ‘cell’ or ‘background’ in this unsupervised manner by OF (dynamic features from image pair (*t* − 1, *t*)), entropy and gradient feature vectors (static features from image at *t*) are generated for each of these training pixels using their local neighborhood of pixels (S.I., Fig S2). These additional feature vectors are then used train and generate a Naïve Bayesian classifier model which is applied to the entire image in a pixel-wise fashion. The information gained from the entropy and gradient feature vectors enables pixels which were left unlabeled in the OF training steps (Fig 2c,d grey pixels) to be classified. The contrast enhanced image (Fig 2b) and model-generated segmentation (Fig 2f, teal pixels) show that the algorithm is able to segment the cell with high fidelity (DIC image/segmented boundary overlay, Fig 2g). Importantly, this labeling, training and classifying procedure occurs recursively on each successive pair of (*t* − 1, *t*) images, enabling the classifier model to adapt to changing backgrounds and phenotypes. By using optical flow to label the highest flow pixels as ‘cells’ and lowest flow pixels as ‘background’, the labeling process has become automated (or ‘self-supervised’) and no manual inputs or training images are needed.

**Fig 2.**
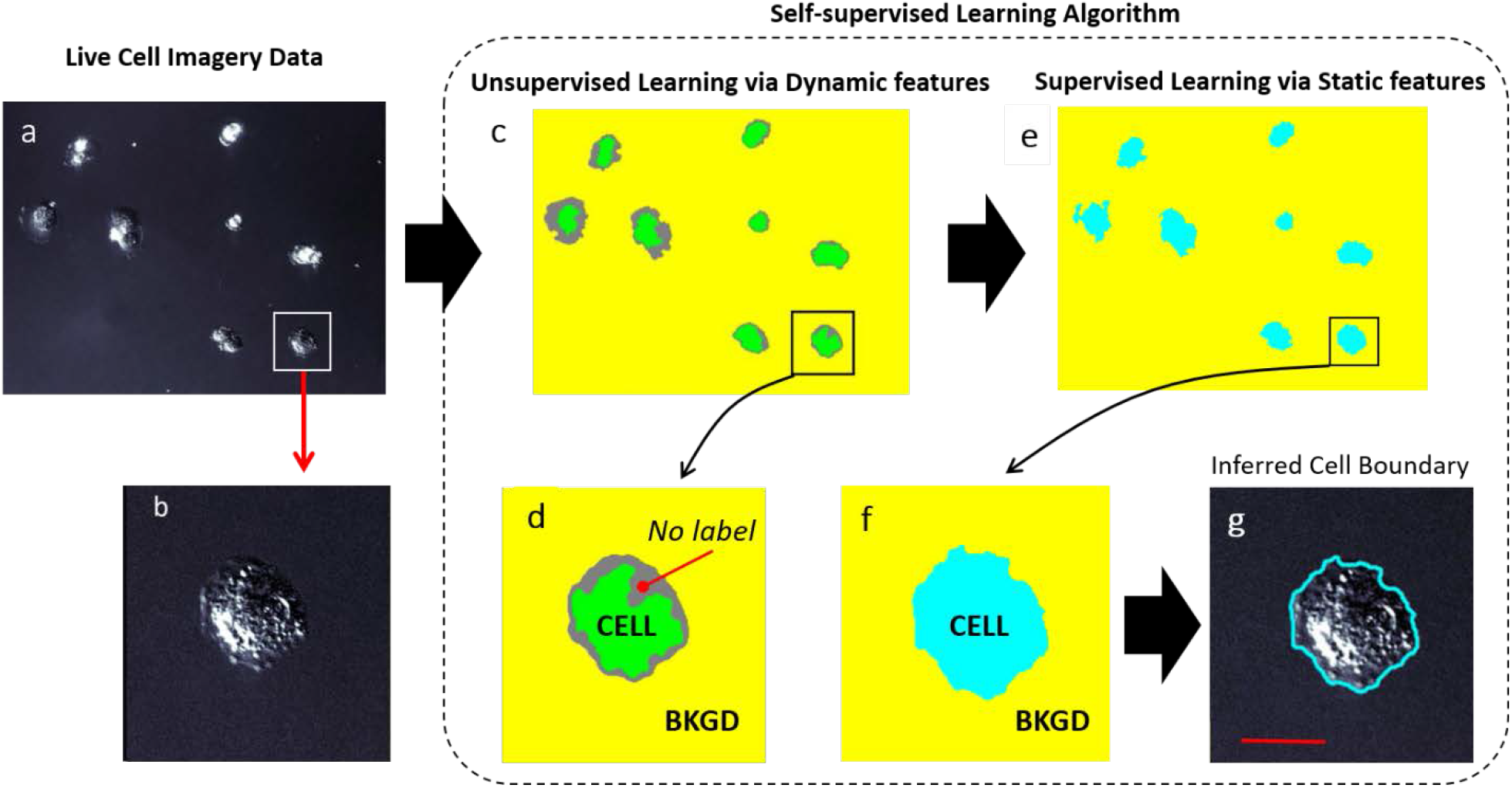
Overview of the automated self-supervised learning algorithm. **a**. The contrast enhanced DIC image of several and **b** a single highlighted MDA-MB-231 cell illustrates the range of intensities inherent within the cells. (20X objective). **c. & d**. Unsupervised learning via OF: high threshold OF is used to select only those pixels exhibiting the highest flow magnitudes and labels them as ‘cell’ (green pixels). Similarly, low threshold OF is used to identify pixels with a much wider range of flow magnitudes than the high flow regime. The lowest flow magnitude pixels are labelled ‘background’ (yellow pixels). Pixels that exhibit OF in between these regimes remain unlabeled (gray pixels). **e. & f**. Supervised learning via self-labeled training data. The self-labeled pixels (green and yellow) are then used to generate static feature vectors, which are in turn used to train the classifier model. **g**. The blue outline is the resulting segmentation which outlines all pixels classified by the OF trained model as ‘cell’ and is also overlaid on the image in **b**. This process is repeated at every time step, thereby using the most recent imagery to update the training data. Scale bar: 25 µm (20X objective, time increment: 300 sec).

For extremely low contrast imagery there can be too few training pixels labeled ‘cell’ for robust segmentation to occur given the initial OF threshold setting. In such cases, the algorithm calculates the entropy associated with ‘cell’ pixels and iteratively reduces the OF threshold until the associated ‘cell’ entropy feature vector is well distinguished from that of the ‘background’ entropy feature vector.

## Results

The Fig 3 imagery shows the generality of this approach and also demonstrates how the self-supervised algorithm additionally automates commonly required manual inputs such as size filtering and hole filling. The segmented cells were processed from imagery acquired from a range of cell types, imaging modalities, magnifications and time increments (S.I. Table S1). The OF algorithm enabled a straightforward approach to automated size filtering which is a common user adjustable parameter in supervised machine learning approaches. To accomplish this, a stand-alone application of OF was applied to the imagery which lacked the added steps of self-tuning and model building described above. While some cell features are missed, this simpler, faster approach was found to be more than precise enough to estimate average cell size and to exclude much smaller objects, thus automating the size filtering process. Because extraneous debris often lacked the motion of the live cells, this debris was also automatically labeled as background by the OF algorithm. Fig 3a and b demonstrate the self-supervised code’s ability to size filter, while also adapting to cell types of differing sizes, by comparing the segmentation of human fibroblasts (10X, phase contrast) to those of the much smaller Dictyostelium amoeboid cells (10X, transmitted light), respectively. Extraneous debris features in the Hs27 imagery (Fig 3a, white arrows) are correctly identified as ‘background’, even though similar in size and intensity to the Dictyostelium cells of Fig 3b. The background inhomogeneities observed in Fig 3a and 3b, which could potentially be mislabeled as ‘cell’, are correctly identified because they remain relatively constant from frame *t* − 1 to frame *t*. The segmentation results of the MDA-MB-231 cells (10X, phase contrast) in Fig 3c illustrates the algorithm’s ability to adapt to a wide range of phenotypes, from rounded Fig 3c(i) to spread Fig 3c(ii), which is enabled without need for user input by continuously retraining the model on consecutive image pairs. The current instantiation of the software does not attempt to separate cells that are touching or close enough to be segmented as a single object. Well-developed approaches such as watershed transforms^23^ and levelset methods^24^ can be employed for such purposes.

**Fig 3.**
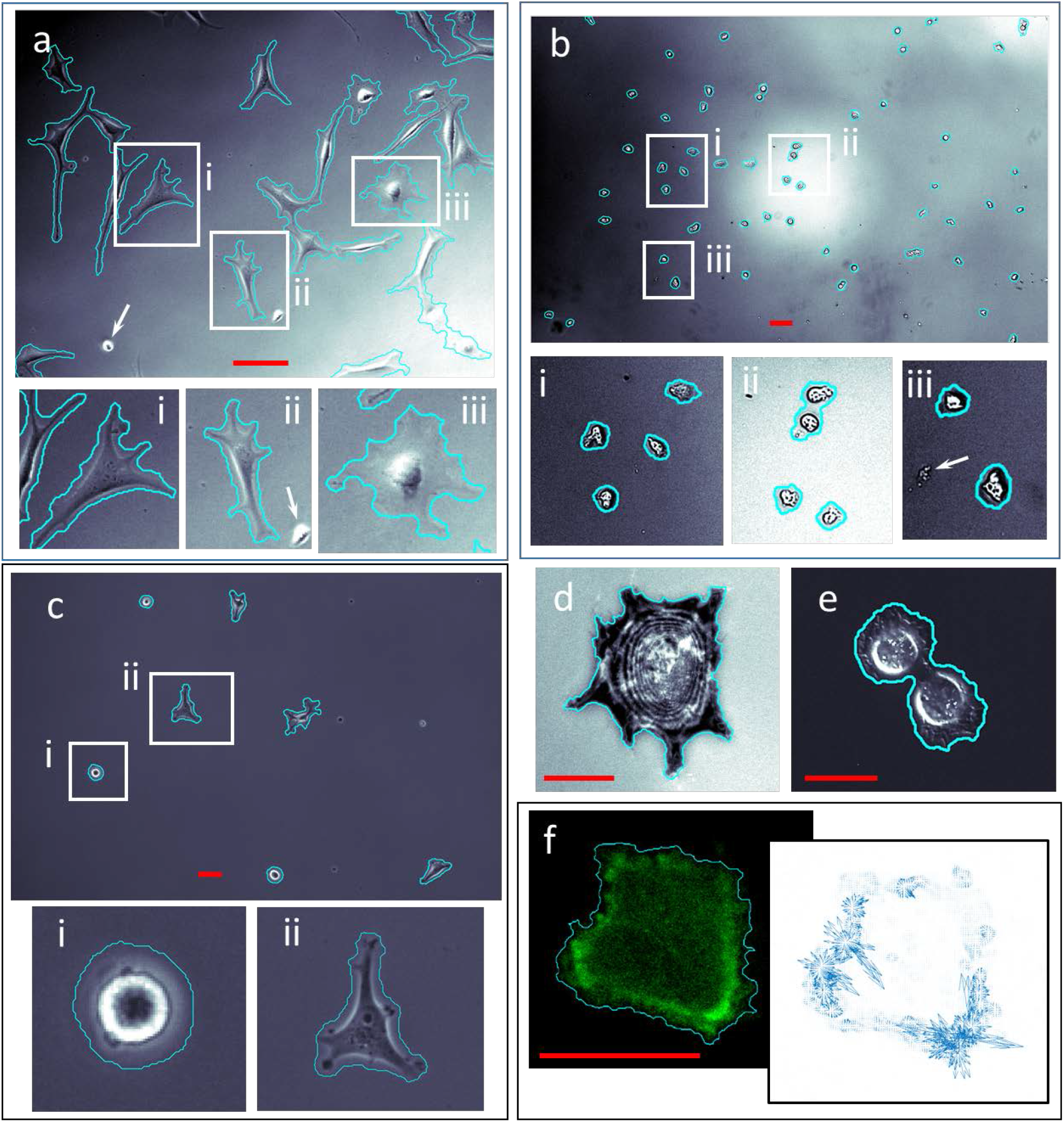
Self-supervised segmentation for a range of cell types, microscope modalities, time resolutions and magnifications. **a**. phase contrast of Hs27 fibroblasts (10X objective, time increment: 1200 sec) **b**. transmitted light of Dictyostelium (10X objective, time increment: 60 sec) **c**. phase contrast of MDA-MB-231 (10X objective, time increment: 600 sec) **d**. IRM image of a single Hs27 cell (40X objective, time increment: 600 sec) **e**. DIC image of MDA-MB-231 cells (20X objective, time increment: 120 sec) **f. f**luorescence image of a single lifeAct (GFP-actin conjugate) transfected A549 cell (pseudo-colored) with the associated optical flow vector plot (100X objective, time increment: 10 sec). Insets i, ii, iii highlight boxed image regions. White arrows point to examples of debris that was correctly labelled ‘background’ due either to lack of motion or automated size filtering. Images have been contrast enhanced to highlight low contrast features and background inhomogeneities. DIC image (**e**) was additionally enhanced with a sharpen filter to highlight interference induced shadowing of cell features. Scale bars: a, b, c: 50 µm; d, e: 25 µm; f: 10 µm.

The algorithm works robustly for a range of optical modalities and magnifications as shown in Figs 3d-f. Figs 3d and 3e are segmentation results from IRM imagery (40X, Hs27 cell) and DIC imagery (20X, MDA-MB-231). As a fluorescence imaging example, a self-supervised segmentation of a GFP-actin labeled A549 cell at 100X magnification is shown in Fig 3f. As an additional option, OF can be applied not only as an algorithm labeling element, but also a measurement tool, as shown in the Fig 3f vector plot. The plotted OF vectors (blue) display the magnitude and direction of the measured GFP labelled actin flow between frames. Such measurements have been shown to be useful for quantifying intracellular protein and calcium signaling dynamics.^25-27^

Hole filling, another often required manual input for model-based and machine learning algorithms, has also been automated by this approach. Common examples of when hole filling input is required include fluorescent labels that do not penetrate the nucleus or, for label-free microscopy modes such as phase contrast, large spread cells in which the algorithm has a difficult time associating the interference enhanced cell edges with the enclosed lamellipodia. We found that motion within cells was ubiquitously detected by OF, regardless of imaging modality or whether imaging the cell membrane, nucleus or cytoplasm. Because motion detection was far more common than not for a given pixel within an area labeled ‘cell’, a fixed morphological blurring tool (circular with a radius of 5 pixels) was found to robustly hole fill regardless of cell type or microscope configuration. The calculated cell area was found to be invariant for a range of blurring tool radii (Fig S2). In all cases, the use of optical flow to identify motion and the 5 pixel radius blurring tool was sufficient to correctly fill in the cell.

By re-training on every pair of consecutive images the self-supervised algorithm remains accurate throughout long-term imaging applications, despite changes in background or cell phenotypes. This allows for a rich behavior of dynamic morphology and migration to readily be collected and analyzed – a key point given the known inter-relationship between cellular shape and function.^2,28,29^ Furthermore, the emerging role that not just cell shape, but cell shape dynamics play in fundamental biological processes is becoming increasing clear.^30^

Fig 4 demonstrates how such quantitative morphological information is readily mined in a long-term imaging application. Fig 4a-c shows the tracking of several MDA-MB-231 cells segmented via the self-supervised approach under 10X phase contrast microscopy on cRGD functionalized gold coverslips.^31^ Fig 4a shows the labeled tracks of the cells’ centroids over the course of 400 minutes, with the corresponding initial and final image shown in Figs 4b,c. The cell associated with track 2 undergoes mitosis at approximately 320 minutes, creating two new tracks (5 and 6) for the daughter cells. Because the self-supervised approach automatically re-trains continuously on consecutive frame pairs, the morphological changes from Fig 4b to Fig 4c are quantified with high fidelity, as can be seen by plotting the segmented boundaries as a function of time (Fig 4d).

**Fig 4.**
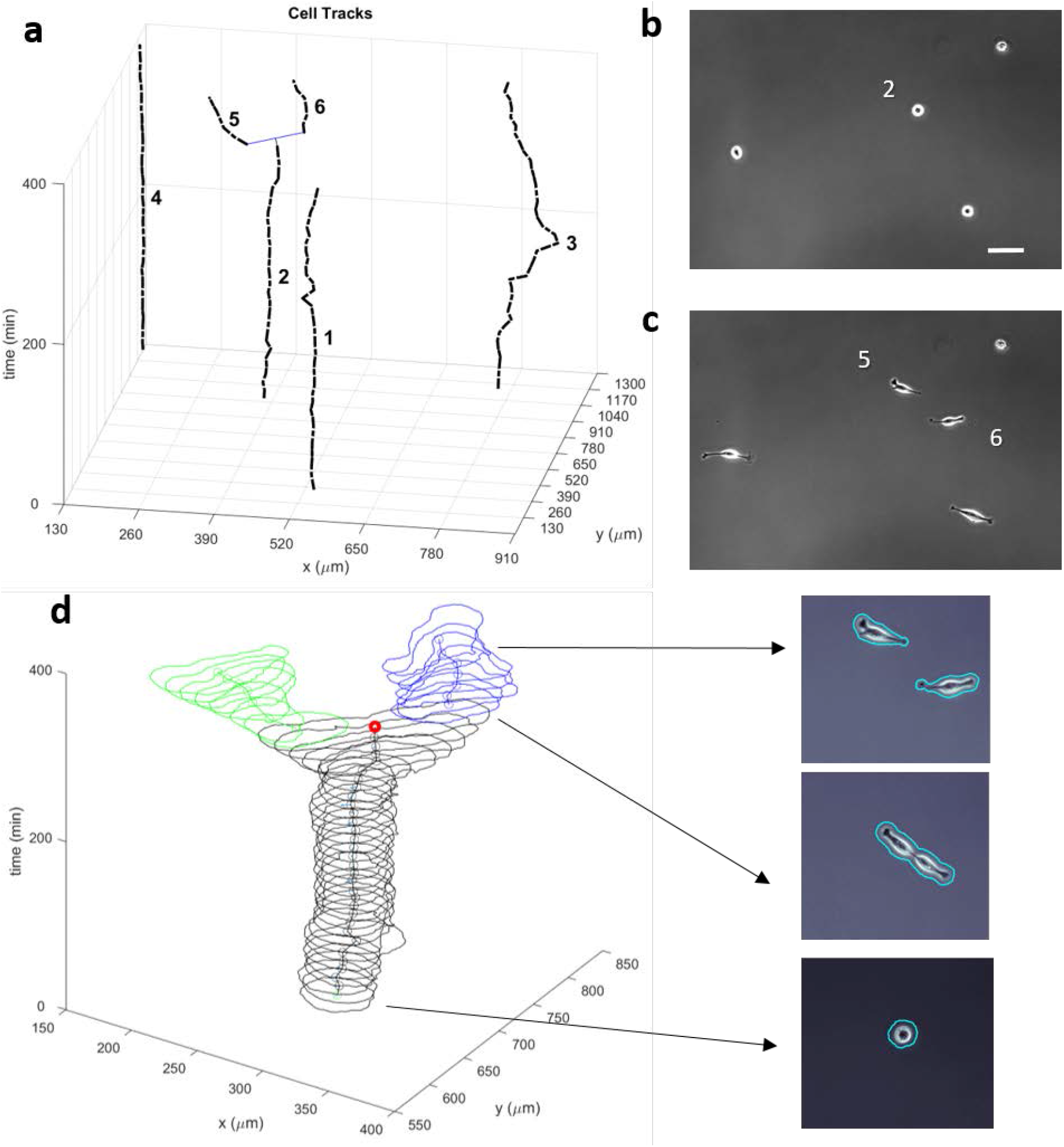
Tracking of MDA-MB-231 cells under 10X phase contrast microscopy and time evolution of cell morphology through mitosis. **a**. The resulting tracks of multiple segmented cells from a single field of view over the course of 400 minutes **b**. corresponding images at times t = 0 min and **c**. 400 min. Track 2 undergoes mitosis resulting in tracks 5 and 6 of the daughter cells (blue line). **d. (left)** Time evolution of segmented morphology of track 2 (black) with the centroid of each shape denoted by an open circle until mitosis, after which the track splits into 5 (green) and 6 (blue), with the cell separation event denoted by a single red open circle. **d. (right)** selected images showing raw data overlapped with the self-supervised segmentation throughout mitosis event. (10X objective, time increment: 600 sec) Scale bar: 100 μm.

## Discussion & Conclusions

There are numerous advantages to this self-supervised machine learning approach. The most obvious is that because the training data is generated by tracking motion, the approach can be used with any live cell imaging microscopy technique, whether labeled or label-free. Also unique is the use of the optical flow labeled pixels to self-supervise the building of a classifier model, which in turn is modular with regards to the incorporated feature vectors. While we have employed only two feature vectors in this current instantiation of the classification code (gradient and entropy), there are many additional image features that can be added based on the application. We have also shown that the incorporation of OF enables the straightforward automation of morphological operations such as size filtering and hole filling, eliminating the need for manually tuning these parameters.

The automation described here is markedly different from machine learning approaches that require user assisted training. The most time consuming aspect of model-based tuning and machine learning approaches is the training process. The process is one of trial and error, requiring retraining if the model’s performance is not deemed adequate. The complete automation of both the training and segmentation algorithms not only saves time but also removes the chances of unconscious bias from entering the training process. Because the training is conducted recursively with each new image, evolutions in phenotype and background structure over extended time periods are accounted for without the need for preprocessing.

The sum of all these advantages is segmentation under a wide range of magnifications, time resolutions, cell types and optical modalities that is both automated and robust. This results in the ability to track cells for hours or days and quantify a range morphological and phenotypic features without the need for user input, thus having broad applicability throughout live cell microscopy. The crux of the introduced self-supervised approach relies upon using the dynamic information embedded in each pixel – motion characterized via optical flow – as an elegant means to self-label cells versus background in time-lapse imagery. While cellular dynamics has long been appreciated as information rich with regards to understanding cell function, our approach demonstrates that it also provides the means for robust segmentation – a foundational step for achieving quantitative and objective live cell analysis.

## Acknowledgements

The authors gratefully acknowledge the Devreotes laboratory of Johns Hopkins University for the *Dictyostelim discoideum* cell line. M.C.R. gratefully acknowledges support from the National Research Council Research Associateship Program and the Jerome and Isabella Karle Distinguished Scholar Fellowship Program. Funding for this project was provided by the Office of Naval Research through the Naval Research Laboratory’s Basic Research Program and by the Biological Technology Office of the Defense Advanced Research Program Agency.

## Author Contributions

Michael C. Robitaille: conceptualization, methodology, investigation, data curation, software, visualization, and writing. Jeff M. Byers: conceptualization, methodology, formal analysis, and software. Joseph A. Christodoulides: Resources, validation, and writing. Marc P. Raphael: conceptualization, funding acquisition, methodology, investigation, software, visualization, and writing.

