## Supplemental Information for "A Self-Supervised Machine Learning Approach for Objective Live Cell Segmentation and Analysis"

#### A. Static Feature Vectors

The self-supervised algorithm allows for automated training of as many static feature vectors as deemed helpful to robustly segmenting the cells from the background. In this work we utilized the entropyfilt(I,nhood) function in MATLAB to calculate the entropy of image I from a surrounding neighborhood of pixels, nhood, which was fixed to a 7 x 7 matrix. In a similar manner, we utilized the imgradient(I,method) function to assign a gradient to a given pixel based on 5 x 5 matrix neighborhood calculated with the 'intermediate' gradient operator method. An example of these feature vector histograms from Dictyostelium time course imagery are shown in Fig S1. Whether a pixel was labeled 'cell' or 'background' was determined by OF.

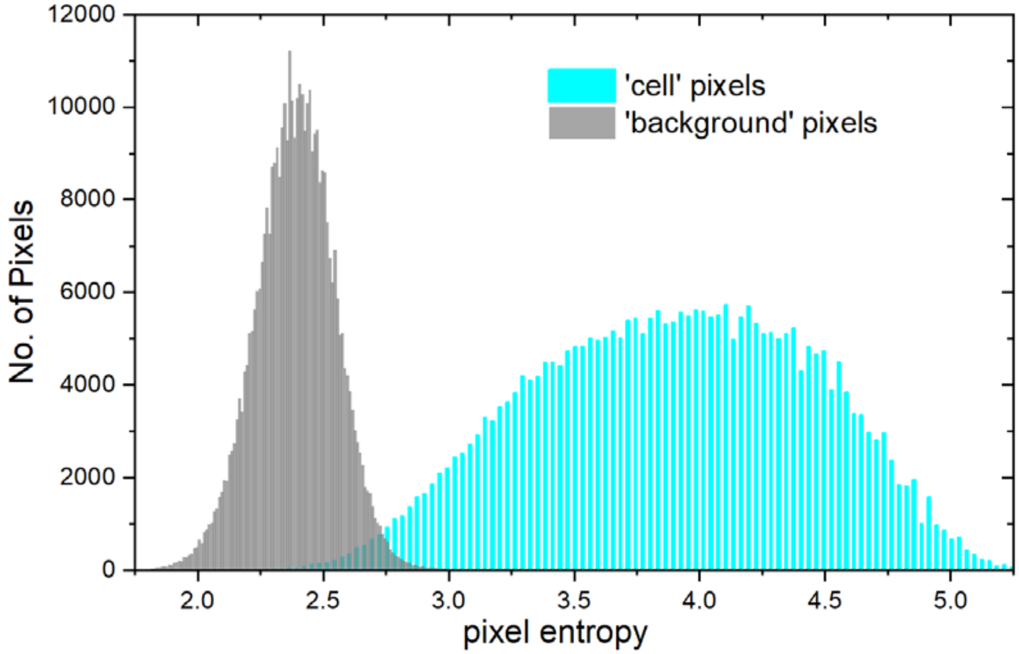

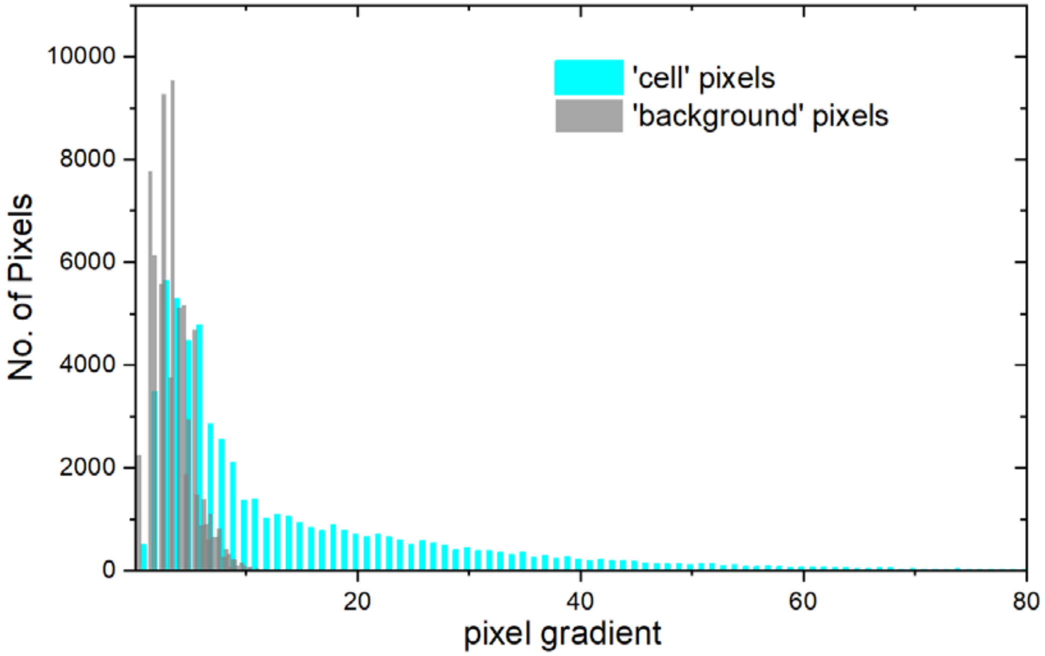


**Fig S1.** (top) Associated entropy feature vector histograms and (bottom) gradient feature vector histograms of the 'cell' and 'background' training pixels used for model training from transmitted light Dictyostelium images (10X objective).

### B. Microscopy Details

| Main Text Figure | Cell Type | Optical Modality | Objective Magnification | Objective Numerical Aperture | Camera | Wait Time [t-1, t] (seconds) |
|---|---|---|---|---|---|---|
| **Figure 2** | MDA-MB-231 (human breast adenocarcinoma) | DIC | 20X | 0.8 (air) | Zeiss Axiocam 702 CMOS | 300 |
| **Figure 3a** | Hs27 (human foreskin, fibroblast) | Phase | 10X | 0.45 (air) | Hamamatsu ORCA R2 CCD | 1200 |
| **Figure 3b** | Dictyostelium Discoideum (amoeboid) | TL | 10X | 0.3 (air) | Zeiss Axiocam 702 CMOS | 60 |
| **Figure 3c** | MDA-MB-231 (human breast adenocarcinoma) | Phase | 10X | 0.3 (air) | Zeiss Axiocam 702 CMOS | 600 |
| **Figure 3d** | Hs27 (human foreskin, fibroblast) | IRM | 40X | 1.4 (oil) | Hamamatsu ORCA R2 CCD | 600 |
| **Figure 3e** | MDA-MB-231 (human breast adenocarcinoma) | DIC | 20X | 0.8 (air) | Zeiss Axiocam 702 CMOS | 120 |
| **Figure 3f** | A549 (human lung adenocarcinoma) | Fluorescence | 100X | 1.46 (oil) | iXon Ultra EMCCD | 10 |
| **Figure 4** | MDA-MB-231 (human breast adenocarcinoma) | Phase | 10X | 0.3 (air) | Zeiss Axiocam 702 CMOS | 600 |

**Table S1** Cell and optical details for imagery in the main text. DIC: Differential Interference Contrast; Phase: Phase Contrast; TL: Transmitted Light illumination; IRM: Interference Reflection Microscopy.

C. Segmented Cell Area as a function of Smoothing Disk Size

Hole filling was accomplished with the imclose(I,SE) function in Matlab in which I is the binary input image with 'cell' pixel values 1 and 'background' pixel values 0, and SE is structural element whose size and shape determines the extent of the blurring. We used a disk SE of radius *r* and by plotting the average segmented cell area in the field of view as a function of r we determined a range in which the area output remained relatively constant across all cell types and optical modalities. **Fig. S2a** shows themean segmented area of Hs27 cells (phase contrast, 10X magnification) versus r. From $1 < r < 11$ the segmented area is stabilized, while above $r = 11$ discrete steps in area are observed as a result of cells being grouped together as shown in the cell segmentation images of **Fig. S2b** for $r = 5$ pixels versus **Fig. S2c** for $r = 20$ pixels. The $r = 5$ pixel value was found to reliably hole fill for all cells and optical modalities in this study.

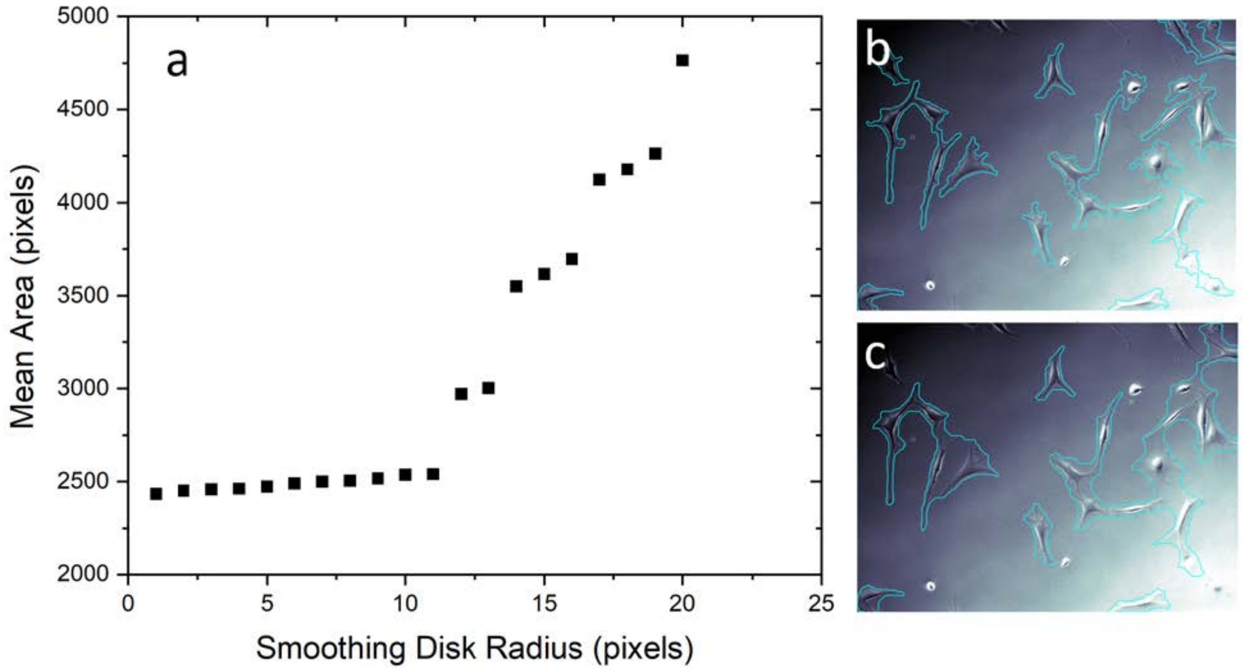


**Fig S2. a.** Mean area of the segmented cells in the field of view as a function of the smoothing disk radius. Segmented images of cells with **b.** $r = 5$ pixels and **c.** $r = 20$ pixels.
